# Dpb11 facilitates the colocalization of Mec1-Ddc2 with its activators on gapped DNA

**DOI:** 10.1101/2025.11.28.691181

**Authors:** Emily C. Beckwitt, Gabriella N. L. Chua, Shixin Liu, Michael E. O’Donnell

## Abstract

The eukaryotic DNA damage and replication stress checkpoint is initiated by activation of the apical kinase complex ATR-ATRIP on RPA-coated ssDNA. In *Saccharomyces cerevisiae*, the Mec1-Ddc2 (hATR-ATRIP) activator and checkpoint mediator Dpb11 (hTopBP1) is recruited to the 9-1-1 checkpoint clamp (another Mec1-Ddc2 activator) at 5’ ss-dsDNA junctions. It remains unclear how Mec1-Ddc2 encounters its activators on damaged DNA due to their differential DNA binding preferences. Using real-time single-molecule imaging, we show that Dpb11 binds to ssDNA directly and localizes to ss-dsDNA junctions in an RPA-dependent manner. Furthermore, Dpb11 recruits Mec1-Ddc2 to ss-dsDNA junctions. Single-molecule force spectroscopy was used to demonstrate that Dpb11 forms bridges on ssDNA, both alone and in the presence of RPA, reducing the end-to-end distance of gapped DNA. These data support a model in which Dpb11 facilitates Mec1-Ddc2 colocalization with its activators directly by recruiting Mec1-Ddc2 to gap junctions and indirectly by decreasing the effective gap length.

## INTRODUCTION

The DNA damage and replication stress checkpoint comprises crucial cellular responses for the maintenance of genome stability and prevention of carcinogenesis^1,2^. This checkpoint can be triggered by single-stranded DNA (ssDNA) resulting from genotoxic challenges during the G1, S, or G2 phase of the cell cycle. Exposed ssDNA is rapidly bound by RPA^3^, which recruits the essential ataxia telangiectasia and Rad3-related (ATR) kinase to the gapped DNA. ATR plays a key role in checkpoint signaling, DNA damage responses, replication fork stability, and control of origin firing^4^. The spatiotemporal mechanisms by which the initial signaling factors of the DNA damage checkpoint pathway are recruited to DNA damage and produce an active signaling complex are poorly understood.

In *Saccharomyces cerevisiae*, Mec1 (homolog of *H. sapiens* ATR) forms a dimer with Ddc2 (homolog of *H. sapiens* ATRIP); this dimer self-associates to form a stable heterotetramer^5-7^. Ddc2 directly interacts with RPA, thereby recruiting Mec1 to ssDNA^6,8-11^. The kinase then needs to be activated, which occurs via interaction with at least one of its three activators: Ddc1^12^, Dpb11^13,14^, and Dna2^15^. These proteins each possess a Mec1-activating domain (MAD) contained in a long unstructured region that stimulates Mec1^16-19^. Ddc1 is one subunit of the heterotrimeric ring called 9-1-1, which is loaded around DNA at a 5’ single-stranded/double-stranded DNA (ss-dsDNA) junction by the pentameric clamp loader Rad24-RFC (homolog of *H. sapiens* Rad17-RFC)^20-23^. The currently accepted model for Mec1 activation in yeast during G2 phase suggests that Ddc1 activates Mec1 both directly via its own MAD and indirectly by recruiting Dpb11^17,24^. During S phase, the checkpoint can also be triggered by the Dna2 nuclease^18,25^.

Dpb11 (homolog of *H. sapiens* TopBP1) is a scaffold protein that can bind multiple phosphorylated proteins via its four BRCA1 C-terminal (BRCT) domains^26,27^ and plays a role in replication initiation and the DNA damage response. The functional role of Dpb11 is governed by its binding partners. Dpb11 activation of Mec1 is highly conserved in eukaryotes^13^. During checkpoint signaling, Dpb11 is able to simultaneously bind phosphorylated Ddc1 and Rad9 (a checkpoint mediator which also binds modified histones^28,29^ and the effector kinase Rad53^30-32^). Interaction between Dpb11 and Ddc1 facilitates recruitment of Dpb11 to gapped DNA. As such, it is generally believed that Dpb11 is, at least in part, dependent on 9-1-1 loading at a 5’ ss-dsDNA junction for cellular activation of Mec1^17^. However, biochemical data demonstrating direct interaction of Dpb11 with various DNA structures or RPA^33^ suggest the possibility that Dpb11 may be recruited to damaged DNA independently of 9-1-1.

Short ssDNA gaps (35 – 200 nt) trigger a weak checkpoint response and longer gaps (900 – 5000 nt) trigger a more robust response by *X. laevis* egg extracts^34,35^. Activation of the global DNA damage checkpoint is highly dependent on the length of ssDNA, with gaps lengths > 1 kb eliciting the most robust response^36^. Processing of double-strand breaks during break-induced replication (BIR) following replication fork collapse generates approximately 1 kb ssDNA^37^. Outside of S phase, 2-6 kb ssDNA are generated during homologous recombination and 10-15 kb ssDNA are generated during BIR^38^. When repair DNA synthesis is overwhelmed, the short gaps of nucleotide excision repair (NER) intermediates can become extended up to several kb and trigger the DNA damage checkpoint^39,40^. Although these long distances are associated with maximal checkpoint activation, they also pose a challenge because the Mec1 activators (Ddc1 and Dpb11) are found at the 5’ junction of a gap and Mec1-Ddc2 is found on RPA at internal gap sites. Therefore, how can Mec1-Ddc2 and the Mec1 activators achieve the proximity required to successfully activate the checkpoint?

In this work, we used combined single-molecule fluorescence and force microscopy^41^ to directly address four major questions: (1) What is the binding specificity of Dpb11 on gapped DNA? (2) How is Mec1-Ddc2 recruited to gapped DNA in the presence of its activator Dpb11? (3) Can Dpb11 alter the structure of gapped DNA, possibly facilitating its connection to Mec1-Ddc2? (4) What is the stoichiometry of Dpb11, a parameter that may help explain the ability of Dpb11 to form multiple protein and DNA interactions? We report that Dpb11 can recognize and bind gapped DNA, even in the absence of 9-1-1. Strikingly, we found that Mec1-Ddc2 binds RPA-ssDNA with no preference for gap junctions, while Dpb11 binds preferentially to ss-dsDNA junctions in the presence of RPA. Perhaps most importantly, we showed that Dpb11 changes the binding specificity of Mec1-Ddc2 and biases it towards ss-dsDNA junctions. This may help to explain a sub-pathway for DNA damage checkpoint activation that does not depend on 9-1-1. Moreover, we found that Dpb11 can bridge distant points on a long DNA molecule in the absence or presence of RPA, stabilizing ssDNA loops that may assist in localizing Mec1-Ddc2 to ss-dsDNA junctions. We have also determined that Dpb11 oligomerizes readily, existing predominantly as a dimer. Our single-molecule studies provide a working model that explains the colocalization of Mec1-Ddc2 and its multiple activators on damaged DNA and reveals additional roles for Dpb11 scaffolding in checkpoint activation.

## RESULTS

To study the recruitment and coordinated assembly of early DNA damage checkpoint factors on damaged DNA, we prepared purified RPA, Mec1-Ddc2, and Dpb11 (**Figure** S1A-B). We first confirmed that our purified proteins were active using a previously reported ensemble phosphorylation assay^12,13^. Mec1 activity was assessed in the absence and presence of activator Dpb11. Dpb11 was indeed able to stimulate Mec1-dependent phosphorylation of Ddc2, Dpb11, and RPA (**Figure** S1C), indicating that each of our purified proteins behave as expected, enabling Dpb11 to activate the Mec1 kinase.

### Dpb11 directly recognizes gapped DNA

Because DNA damage and replication stress checkpoint signaling is broadly initiated by persistent, long regions of ssDNA, we chose to investigate Dpb11 and Mec1-Ddc2 recruitment to a DNA molecule containing a 3 kb single-stranded gap. Protein binding events were followed in real time using a Lumicks C-Trap that combines dual-trap optical tweezers and three-color confocal fluorescent microscopy^42,43^ (**Figure** 1A). Biotinylated 18,851 bp DNA containing two site-specific nicks (at positions 5,982 and 9,018 bp) was tethered between two optically trapped streptavidin-coated beads in a microfluidic flow cell. The DNA was stretched under flow to melt and release the fragment between the nicks, leaving a 3,036 nt ssDNA gap sandwiched between two long dsDNA arms (**Figure** S2A). Successful generation of the gapped DNA was confirmed by both the force-extension profile of the DNA (**Figure** S2B) and the specific staining of YO-PRO-1 dye to the dsDNA regions (**Figure** S2C). Kymographs were recorded under low tension (<20 pN) and in the absence of flow. DNA binding experiments were performed in a flow cell chamber that was free of DNA stains and contained purified *S. cerevisiae* proteins.

**Figure 1.**
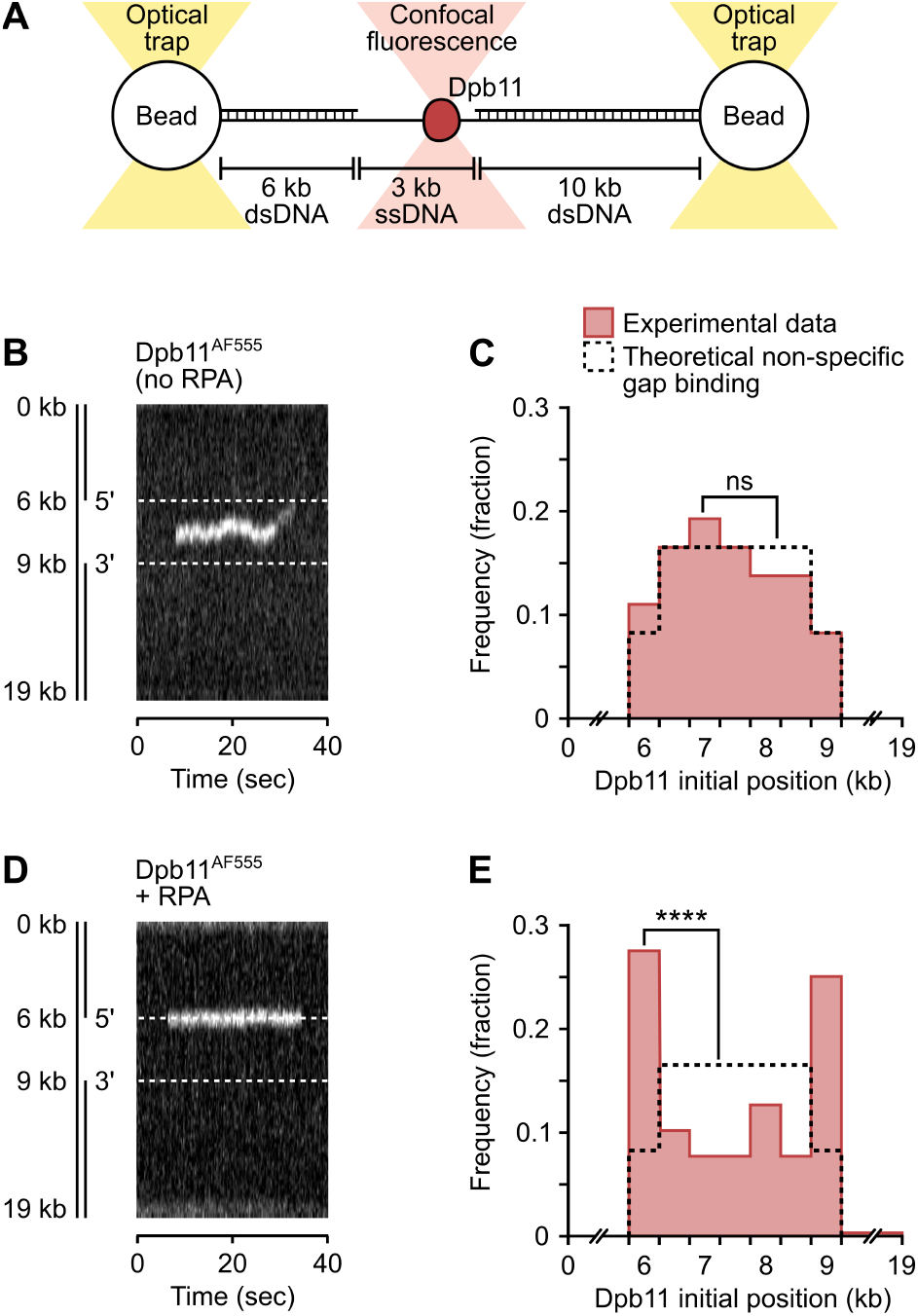
Dpb11 binds preferentially to 3’ and 5’ ss-dsDNA junctions in the presence of RPA. (A) Experimental set-up for kymograph collection. (B and C) Single-molecule imaging of Dpb11^AF555^ on 19 kb DNA with a 3 kb single-stranded gap (6–9 kb) in the absence of RPA. Representative kymograph (B) and histogram of distribution of initial Dpb11 binding positions (C) are shown. Binding positions were measured as the distance from Dpb11 to the end of the DNA (0 kb). n = 36 binding events. (D and E) Single-molecule imaging of Dpb11^AF555^ on gapped DNA in the presence of unlabeled RPA. Representative kymograph (D) and histogram of distribution of initial Dpb11 binding positions (E) are shown. n = 42 binding events. Distribution data were compared to the theoretical non-specific ssDNA (C) or non-specific RPA-ssDNA (E) binding distribution by χ^2^ test for goodness of fit. ns, *p* = 0.99; **** *p* = 4.0×10^-6^.

We first asked if Dpb11 could recognize the gapped DNA on its own. Dpb11 was covalently labeled at the N-terminal amine with fluorophore AF555 (Dpb11^AF555^) or LD655 (Dpb11^LD655^) (**Figure** S1A). Dpb11 samples labeled with either fluorophore exhibited similar binding behavior and were thus considered equivalent for analyses (**Figure** S2D). Dpb11 was observed to readily bind the gapped DNA (**Figure** 1B, **Figure** S2E). Importantly, all Dpb11 particles observed on DNA were located within the 3 kb ssDNA gap; none (0%) were detected on the duplex regions. This suggests that Dpb11 has very low affinity for dsDNA compared to ssDNA. Furthermore, plotting the frequency of initial binding position of individual Dpb11 along the DNA molecule reveals no specificity for particular ssDNA positions or ss-dsDNA junctions (**Figure** 1C). The distribution of binding positions matches the theoretical distribution for a protein with sequence-independent affinity for ssDNA, but not for dsDNA or ss-dsDNA junctions (hereafter referred to as “non-specific ssDNA-binding,” see methods).

### Dpb11 binds specifically to DNA gap junctions in the presence of RPA

Because RPA coats cellular ssDNA, we next examined the binding behavior of Dpb11^AF555^ on the 3 kb-gapped DNA in the presence of saturating amounts of RPA (**Figure** 1D, **Figure** S2F). Under these conditions, the binding positions of Dpb11 were significantly different than the non-specific ssDNA-binding distribution observed in the absence of RPA (**Figure** 1E). We observed a clear enrichment at both the 5’ and 3’ ss-dsDNA junctions. Of all the Dpb11 binding events, 53% occurred at a gap junction. Specifically, 28% and 25% occurred at the 5’ junction and 3’ junction, respectively. These values are significantly higher than predicted by the non-specific model (8.3% at either junction), indicating specificity for junctions in the presence of RPA.

Additionally, RPA increases the frequency of stationary Dpb11 particles (**Figure** S2G). In the absence of RPA, 44% of Dpb11 particles exhibited one-dimensional diffusion (i.e. were mobile on DNA) and 56% remained stationary. In the presence of RPA, the vast majority (95%) of Dpb11 was stationary. Thus, RPA appears to limit the diffusive behavior of Dpb11 on ssDNA.

### Mec1-Ddc2 binds RPA-ssDNA

Mec1-Ddc2 is known to be recruited to RPA-bound ssDNA via Ddc2-RPA interactions^6,10,44^. We sought to confirm this behavior using our C-trap setup. Mec1-Ddc2 was covalently labeled at the N-terminal amine with fluorophore AF647 (Mec1-Ddc2^AF647^), LD555 (Mec1-Ddc2^LD555^), or LD655 (Mec1-Ddc2^LD655^) (**Figure** S1B). Mec1-Ddc2 samples labeled with each fluorophore exhibited similar behavior in terms of both kinase activity (**Figure** S1C) and DNA binding (**Figure** S3A). They were thus considered equivalent for analyses. In the presence of saturating amounts of RPA, Mec1-Ddc2 was observed to bind the gapped DNA, but not duplex DNA (**Figure** 2A, **Figure** S3B). The distribution of Mec1-Ddc2 binding positions closely matches the expected non-specific ssDNA-binding model (**Figure** 2B).

**Figure 2.**
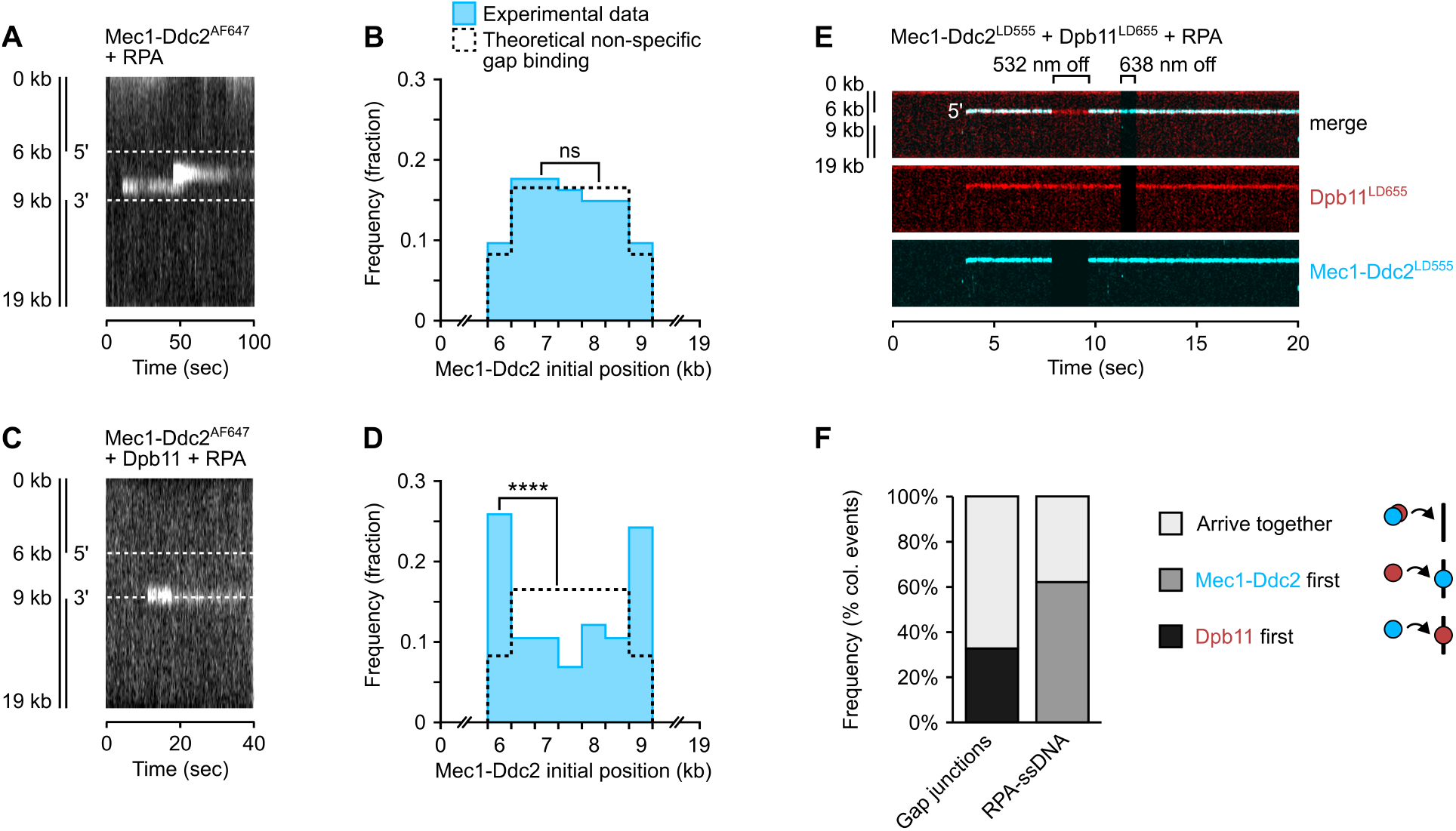
Dpb11 directs Mec1-Ddc2 to ss-dsDNA junctions. (A and B) Single-molecule imaging of Mec1-Ddc2^AF647^ on 19 kb DNA with a 3 kb single-stranded gap (6–9 kb) in the presence of unlabeled RPA. Representative kymograph (A) and histogram of distribution of initial Mec1-Ddc2 binding positions (B) are shown. n = 74 binding events. (C and D) Single-molecule imaging of Mec1-Ddc2^AF647^ on gapped DNA in the presence of unlabeled RPA and unlabeled Dpb11. Representative kymograph (C) and histogram of distribution of initial Mec1-Ddc2 binding positions (D) are shown. n = 58 binding events. (E) Representative kymograph of colocalized Dpb11^LD655^ and Mec1-Ddc2^LD555^ binding to the 5’ junction of 3 kb-gapped DNA. The 532 nm and 638 nm excitation lasers were briefly turned off to confirm specificity of the LD555 and LD655 fluorophores. The merged image is shown above the separated color channels; white indicates colocalized signals. (F) Relative frequencies of each type of colocalization pathway at different positions on gapped DNA. Possible pathways for colocalization of Mec1-Ddc2 (blue) and Dpb11 (red): both proteins interact in solution and bind DNA together (as shown in E), Mec1-Ddc2 binds DNA first and then recruits Dpb11, or Dpb11 binds DNA first and then recruits Mec1-Ddc2. n = 11 colocalization events. Distribution data were compared to the theoretical non-specific RPA-ssDNA-binding distribution by χ^2^ test for goodness of fit. ns, *p* = 1.0; **** *p* = 1.9×10^-8^.

### Dpb11 recruits Mec1-Ddc2 to gap junctions

To determine if Dpb11 has an influence on the location of Mec1-Ddc2, we next observed Mec1-Ddc2 binding to the 3 kb-gapped DNA in the presence of saturating unlabeled RPA and excess unlabeled Dpb11 (**Figure** 2C, **Figure** S3C). In the presence of Dpb11, the distribution of Mec1-Ddc2 binding positions is changed drastically, exhibiting a marked enrichment at both the 3’ and 5’ ss-dsDNA junctions and significantly deviating from the theoretical non-specific ssDNA-binding distribution (**Figure** 2D). Interestingly, the distribution of Mec1-Ddc2 resembles that observed for Dpb11 binding to RPA-coated gapped DNA (compare **Figure** 1E and **Figure** 2D). Like Dpb11, half of all Mec1-Ddc2 binding events occurred at a ss-dsDNA junction.

In the absence of Dpb11, 89% of the observed Mec1-Ddc2 particles were stationary on DNA. In the presence of Dpb11, 83% of Mec1-Ddc2 particles were stationary. These data suggest that, under either condition, Mec1-Ddc2 performs a primarily three-dimensional search in solution for its target. This is consistent with a previous report in human cells^45^.

To directly visualize the interaction between Mec1-Ddc2 and Dpb11 on gapped DNA, we next repeated the experiment with fluorescent Mec1-Ddc2 (Mec1-Ddc2^LD555^), fluorescent Dpb11 (Dpb11^LD655^), and excess unlabeled Dpb11 and unlabeled RPA. We observed colocalization events of Mec1-Ddc2^LD555^ and Dpb11^LD655^ (**Figure** 2E). Colocalization was the result of three distinct pathways: (1) Dpb11 and Mec1-Ddc2 interact in solution and bind DNA together, (2) recruitment of Dpb11 to DNA-bound Mec1-Ddc2, or (3) recruitment of Mec1-Ddc2 to DNA-bound Dpb11. Experimentally, these pathways were categorized based on the order that fluorescent signals for Dpb11^LD655^ Mec1-Ddc2^LD555^ were observed on a given DNA site. We observed a combination of all three pathways on gapped DNA (**Figure** 2F). Interaction between Dpb11 and Mec1-Ddc2 in solution led to colocalization at all positions. Mec1-Ddc2 that is already bound to RPA-ssDNA appears to recruit Dpb11 to internal ssDNA sites, but not to a gap junction. Most interestingly, Dpb11 recruits Mec1-Ddc2 to a gap junction (**Figure** 2F), but not to internal sites. These data support the notion that Mec1-Ddc2 binding to gap junctions is mediated by its interaction with Dpb11, although we acknowledge the possibility of alternative pathways due to the small sample size of colocalized dual-color Mec1-Ddc2 and Dpb11 (n = 11).

### Dpb11 can engage multiple DNA sites and establish DNA loops

Dpb11 is a scaffold protein that can simultaneously bind multiple partners via its four BRCT domains^26,46^. This, combined with our DNA-binding data, led us to ask if Dpb11 can bind DNA in two places at once. To this end, we used single-molecule force spectroscopy to conduct DNA pulling tests (**Figure** 3A), similar to those previously reported^47,48^. This pulling procedure identifies DNA loops bridged by Dpb11, because the presence of loops decreases the effective length of the DNA between the beads. Increasing tension disrupts the bridges, releasing the loops and resulting in a sudden drop in force.

**Figure 3.**
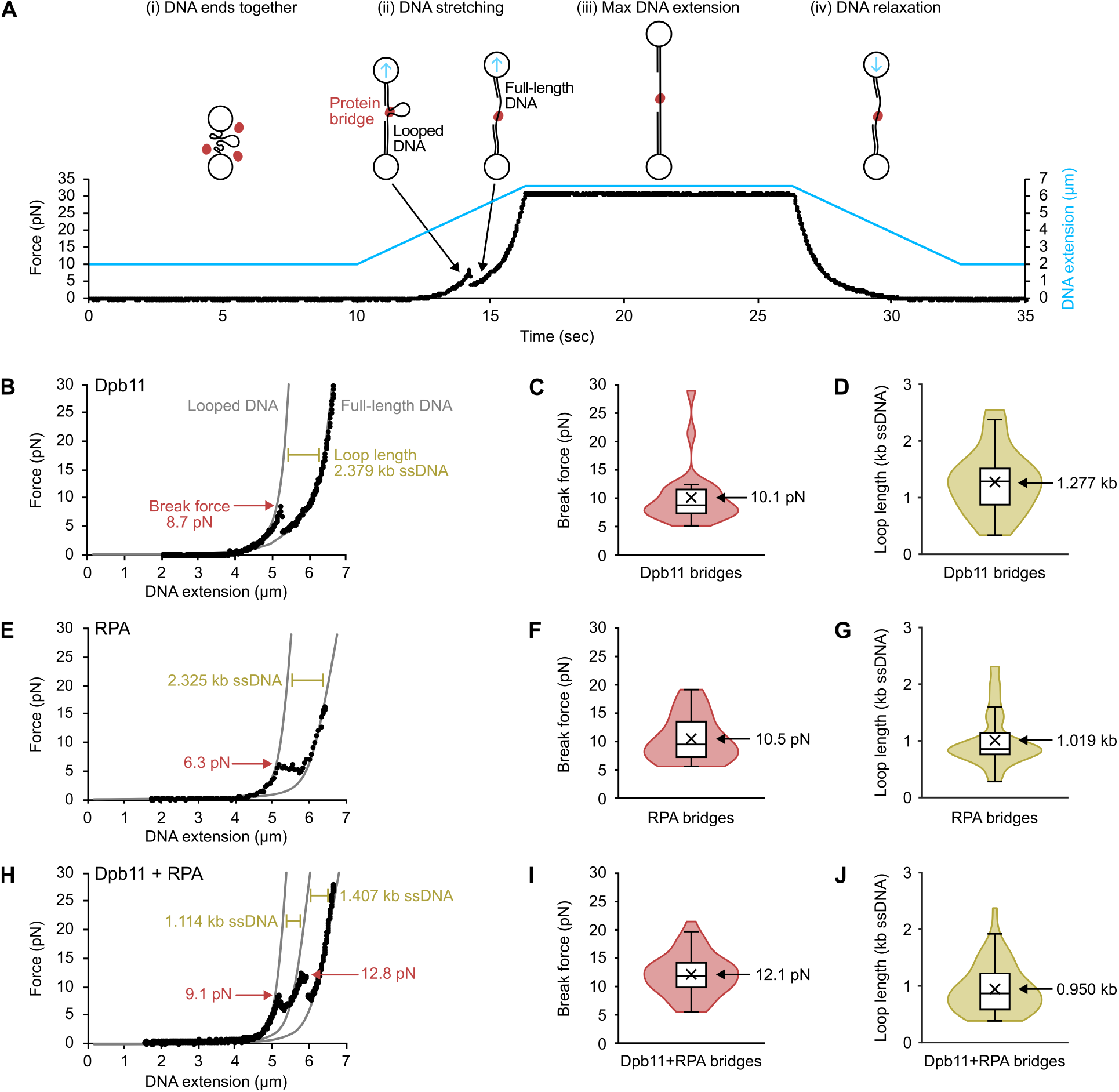
Dpb11 and RPA form bridges on ssDNA. (A) Typical DNA pulling experiment is shown with corresponding force data (left axis) and DNA extension data (right axis) as a function of time. (B–D) DNA pulling tests of Dpb11 and 3 kb-gapped DNA (15.815 kb dsDNA + 3.036 kb ssDNA). (B) Example force-extension plot. Black circles are experimental data. Break force (red) is the maximum force before loop is released. Loop length (olive) is the difference in ssDNA determined by fitting the data before and after the “break” to the appropriate DNA models (gray lines). Summary of Dpb11 bridge break forces (C) and loop lengths (D) are shown. n = 25 bridges. (E–G) DNA pulling tests of RPA and 3 kb-gapped DNA. An example force-extension plot (E), summary of RPA bridge break forces (F) and loop lengths (G) are shown. n = 26 bridges. (H–J) DNA pulling tests of Dpb11 + RPA and 3 kb-gapped DNA. An example force-extension plot (H), summary of Dpb11 + RPA break forces (I) and loop lengths (J) are shown. n = 53 bridges. Summary data (C, D, F, G, I, J) are represented by truncated violin plots with box-and-whisker (Tukey) overlay. The sample means are labeled (×).

The force at which each protein bridge was disrupted was defined as the maximum force before an observed change in DNA length, referred to here as the “break force” (**Figure** 3B). As a negative control, gapped DNA was first tested in buffer. Only 17% of all pulls resulted in a noticeable change in DNA length (**Figure** S4A). These events were likely the result of secondary structure within the single-stranded region of the gapped DNA. Of these, the break force was 3.5 ± 0.8 pN (mean ± s.d.). Thus, we considered any break forces below 5 pN as non-specific background and excluded them from subsequent analyses.

Prevalent bridging (95% of all pulls) was detected on 3 kb-gapped DNA in the presence of only Dpb11 (**Figure** 3B, **Figure** S4B). Dpb11 bridges were disrupted by 10.1 ± 5.1 pN (mean ± s.d.) of force (**Figure** 3C), much higher than the break force observed without Dpb11. The amount of DNA contained in each loop was determined by fitting a portion of the experimental data before and after the break to the appropriate DNA model and solving for the number of ssDNA residues; the difference between the two values provides the “loop length” (**Figure** 3B). DNA loops contained 1,277 ± 565 nt ssDNA (**Figure** 3D). Dpb11 bridges were not formed on the same 18,851 bp DNA substrate without the gap (i.e. 100% dsDNA), indicating that Dpb11 bridges are dependent on ssDNA (**Figure** S4C). This is further supported by the observed low affinity of Dpb11 for dsDNA (**Figure** 1). Thus, we conclude that all bridges were most likely formed within the 3 kb ssDNA region.

### Dpb11 and RPA form multiple bridges on gapped DNA

We next asked if Dpb11-mediated DNA bridges would also form in the presence of RPA. RPA alone can form bridges on gapped DNA (**Figure** 3E, **Figure** S4D) but not on dsDNA (**Figure** S4E). These bridges were disrupted by 10.5 ± 4.9 pN of force (**Figure** 3F) and each loop contained 1,019 ± 483 nt ssDNA (**Figure** 3G). DNA pulling tests conducted in the presence of both RPA and Dpb11 also resulted in bridge/loop formation on gapped DNA (**Figure** 3H, **Figure** S5A). The bridges generated in the presence of the two proteins were disrupted by 12.1 ± 3.7 pN of force (**Figure** 3I) and contained 950 ± 435 nt of ssDNA (**Figure** 3J). Pulling tests conducted with either 200 nM Dpb11 or 100 nM RPA separately resulted primarily in formation of a single bridge. When Dpb11 and RPA were present together at the same concentrations, a greater number of bridges were observed on each DNA molecule (**Figure** S5B), indicating that RPA bridges do not inhibit the formation of Dpb11 bridges (and vice versa). Under these conditions, the effective length of the ssDNA region was reduced from 3,036 nt to 848 ± 565 nt (**Figure** S5C). The effective length of the ssDNA region was significantly reduced in all conditions (**Figure** S5C), demonstrating that Dpb11 and/or RPA bridge-formation can reduce the overall distance between gap junctions.

### Dpb11 oligomerizes

Finally, the coincident abilities of Dpb11 to bind DNA junctions, recruit Mec1-Ddc2, and bridge ssDNA, led us to investigate Dpb11 stoichiometry. To address the question of Dpb11 oligomerization, we used mass photometry, which measures the mass of single particles based on light-scattering of a label-free sample at a water-glass interface^49^. Dpb11 in 250 mM NaCl buffer exists as a mixture of monomer (35%) and dimer (64%) at 50 nM (**Figure** 4A). Increasing the Dpb11 concentration to 150 nM shifts the equilibrium to 98% dimer (**Figure** 4B). The emergence of a shoulder at higher molecular weights suggests a small number of trimers or tetramers were present as well. DNA pulling tests and Mec1-Ddc2 recruitment studies above were performed using 200 nM Dpb11. Therefore, the Dpb11 was likely oligomerized in these experiments.

**Figure 4.**
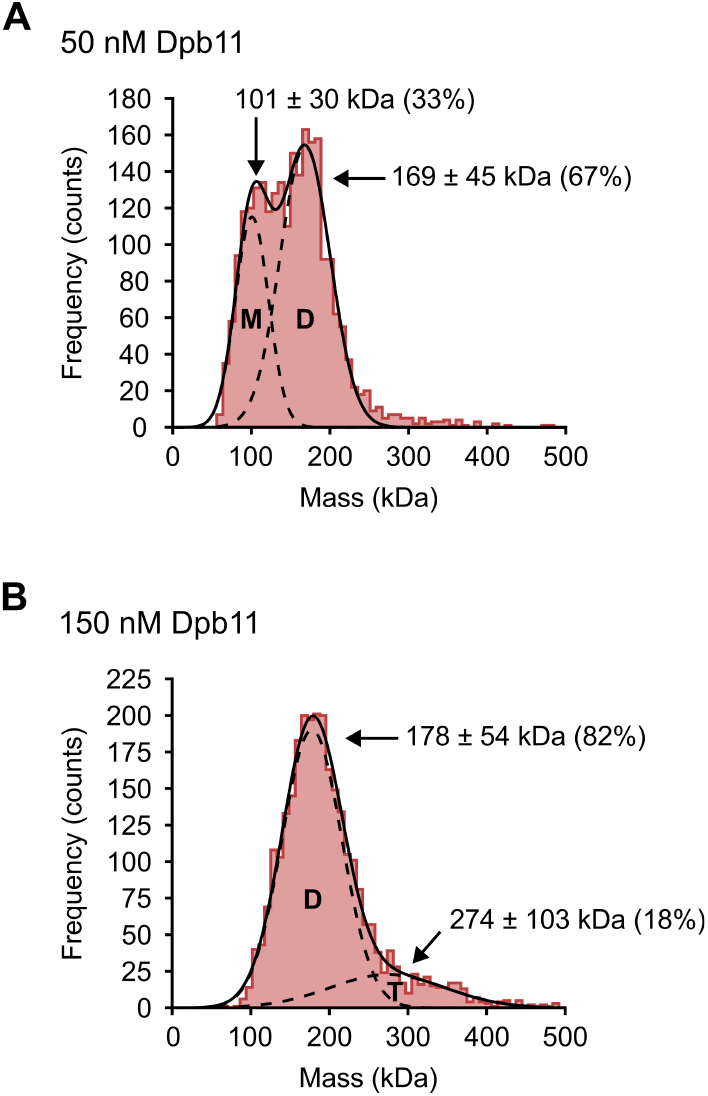
Dpb11 oligomerizes. Dpb11 at 50 nM (A) or 150 nM (B) was analyzed by mass photometry. Molecular weights of individual particles were calculated from raw data using a standard curve of known proteins. Histogram shows mass distribution fitted with a double gaussian (solid black line). Individual gaussians (dashed line) are labeled μ ± σ (% of counts). Most likely oligomeric status of each peak is indicated: M (monomer, 87 kDa), D (dimer, 174 kDa), T (trimer, 271 kDa, or tetramer, 348 kDa).

## DISCUSSION

In the current report, we present a single-molecule study of the coordinated recruitment of two key factors in the DNA damage checkpoint: Dpb11 and Mec1-Ddc2. Previous studies have concluded that Mec1 is recruited to gapped DNA independently from its activators^50,51^. Mec1-Ddc2 binds to RPA on ssDNA^10^, while Dpb11 binds phosphorylated 9-1-1 at a 5’ ss-dsDNA gap junction^24^. The distinct binding sites of kinase and activators present a fundamental problem for checkpoint activation, as they can occur more than 1 kb apart on damaged DNA (**Figure** 5A). How does the cell overcome the potential distance between Mec1 and its activators to trigger the DNA damage checkpoint? We propose a model where Dpb11 has an important gap-recognition function that serves to bring Mec1 within the activation radius of 9-1-1 and Dpb11 in two ways: (1) Dpb11 decreases the effective length of long ssDNA gaps by stabilizing loops, and (2) Dpb11 targets Mec1-Ddc2 to gap junctions where its activators are found. As such, Dpb11’s affinity for gapped DNA is central to its checkpoint activation function (**Figure** 5B).

**Figure 5.**
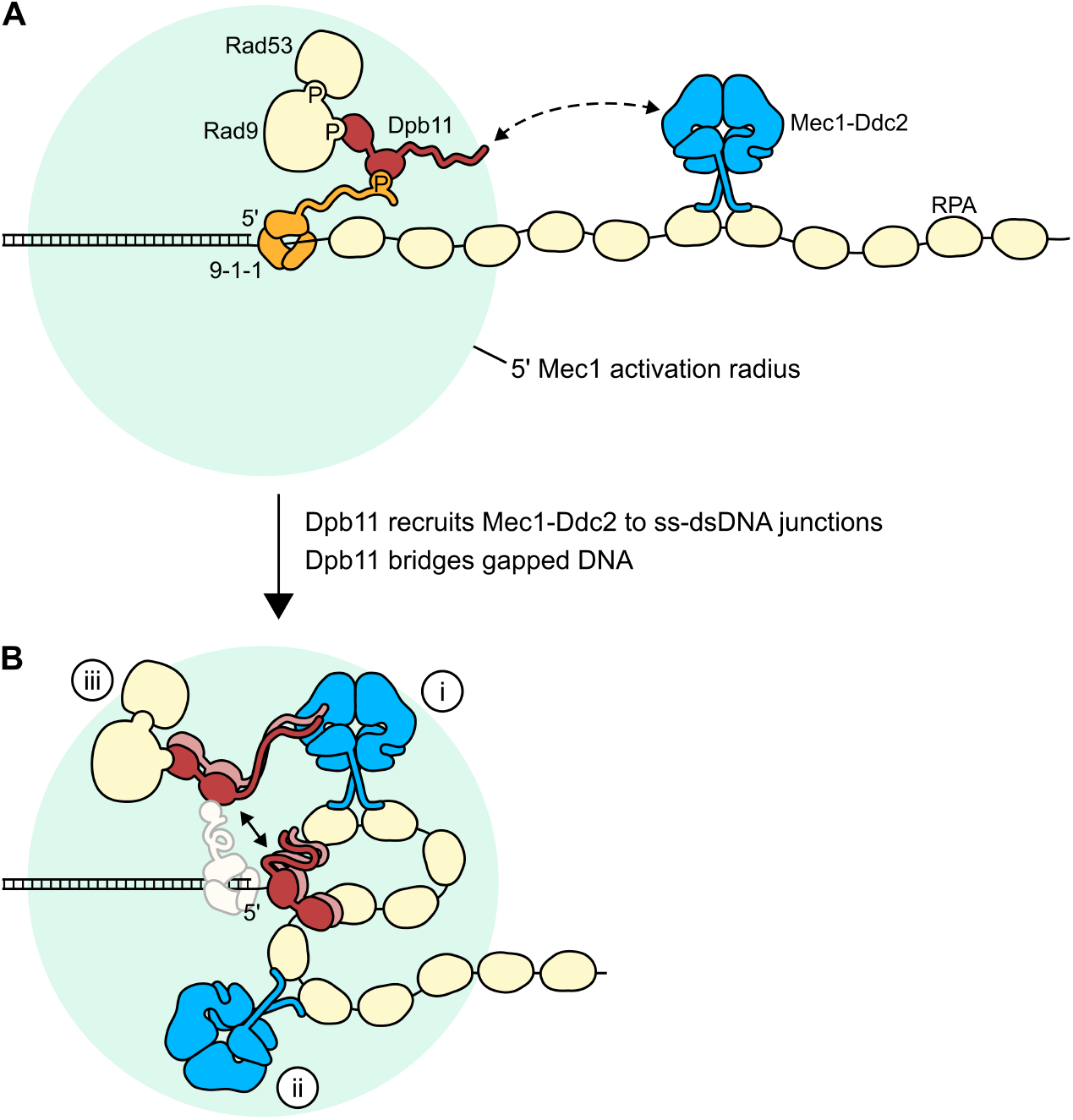
Working model for Mec1-Ddc2 and Dpb11 assembly on gapped DNA. (A) Canonical early steps of DNA damage and replication stress checkpoint signaling on a long ssDNA gap. RPA (pale yellow) binds to ssDNA and Mec1-Ddc2 (blue) binds to RPA. The Mec1 activators Ddc1, as part of the 9-1-1 heterotrimer (orange), and Dpb11 (red) are shown at the 5’ gap junction. The disordered C-terminal tails of Ddc1 and Dpb11 make up the activation radius (pale green circle). The gap-binding functions of Dpb11 identified in the current study help bring Mec1-Ddc2 into the activation radius. (B) Coordinated protein and DNA-binding functions of oligomeric Dpb11 bring Mec1-Ddc2 into the checkpoint activation radius and facilitate assembly of early checkpoint factors. See panel A for protein labels. (i) Mec1-Ddc2, directly recruited to a 5’ ss-dsDNA junction by Dpb11, is activated by Dpb11 or 9-1-1. In S-phase, this may happen in the absence of 9-1-1. (ii) Dpb11/RPA-bridged ssDNA loops bring junction-tethered activators and distant Mec1-Ddc2 complexes into proximity, enabling signal amplification across the length of ssDNA gaps. (iii) Dpb11 scaffolding of Rad9/Rad53 with rest of the assembly facilitates Rad53 activation by Mec1 and checkpoint signal transduction. Illustrated binding poses are hypothetical.

### 9-1-1-independent recruitment of Dpb11 to gapped DNA

A previous study using short oligonucleotides reported that Dpb11 interacts directly with both DNA and RPA^33^. Interactions of TopBP1 (vertebrate homolog of yeast Dpb11) with DNA and RPA also remain unclear as conflicting studies report either that TopBP1 binds ssDNA but not RPA^11,52^ or that it binds RPA but not ssDNA^53^. Here, we directly visualize Dpb11 physically interacting with a 19 kb DNA substrate containing a 3 kb ssDNA gap. Single-molecule imaging along these long distances enables observation of checkpoint signaling aspects that were not detected in the previous studies. In the absence of RPA, we provide definitive evidence that Dpb11 exclusively binds to ssDNA and is not detectable on dsDNA. Dpb11 may bind ssDNA through its BRCT4 domain and C-terminus^54^ (**Figure** S6A). Pairs of BRCT domains canonically bind phospho-proteins, but single BRCT domains have also been shown to bind DNA, as in replication factor C (RFC)^55^. Interestingly, in the presence of RPA, Dpb11 binding is significantly enriched at the ss-dsDNA junctions of gapped DNA. While we observe that Dpb11 has comparable specificity for both the 5’ and 3’ junctions in our minimal *in vitro* system, it is possible that the presence of other proteins, like DNA polymerase at the 3’ junction, may affect this distribution.

### Dpb11 oligomerization

The scaffolding capacity of Dpb11 is at the core of its ability to bring Mec1 into the checkpoint activation radius. Dpb11 can both tether Mec1-Ddc2 to gap junctions and bridge discrete DNA sites. We report that Dpb11 readily dimerizes in solution and may form higher order oligomers at high concentrations. This is consistent with previous reports on vertebrate TopBP1, which likely functions as an oligomer for ATR activation^56-59^. *X. laevis* TopBP1 self-associates via homotypic interactions between the BRCT1+2 domains and heterotypic interactions between the BRCT1+2 and BRCT4+5 domains^59^ (**Figure** S6B). These interactions are direct, do not require any post-translational modifications, and do not interfere with phospho-protein binding activity; they also enable checkpoint activation^59^. Based on protein homology, we expect that Dpb11 dimers contain a BRCT1+2–BRCT1+2 interface and a BRCT1+2–BRCT3+4 interface. Additional homotypic interactions between the vertebrate BRCT7+8 domains have been reported^60^ but these have no known homology with yeast Dpb11 (**Figure** S6B). The validation of oligomer interfaces in yeast warrants further investigation. It is understood that monomeric Dpb11 is able to bridge binding partners that occupy different BRCT domains. Oligomeric Dpb11 can also bridge binding partners that occupy the same positions, as evidenced here by its ability to simultaneously bind discrete ssDNA sites. Additionally, Dpb11 interaction with Mec1-Ddc2 and DNA both involve the C-terminal tail of Dpb11^13,14,33^ (**Figure** S6A). As such, oligomerization may also explain our observation that Dpb11 can simultaneously bind Mec1-Ddc2 and DNA, thereby recruiting Mec1-Ddc2 to gapped DNA. Dpb11 oligomerization and Dpb11-induced DNA looping may also serve a function outside of checkpoint signaling, such as the reported stimulation of ssDNA annealing in the context of origin firing^61^.

### Dpb11 indirectly positions Mec1 within the activation radius: bridges on ssDNA

The discovery that Dpb11 can stabilize long loops of ssDNA has important implications for checkpoint signaling. RPA bridges have been reported on unrestrained ssDNA in a previous study^62^. We demonstrate that both Dpb11 and RPA can form bridges across long regions, separated by an average of 1 kb relaxed ssDNA. These large loops are most likely formed and stabilized by protein bridges and effectively reduce the end-to-end distance of gapped DNA. We postulate that the observation of complete junction-to-junction looping was likely obstructed by technical limitations, such as the exclusion of all break forces below 5 pN or the requirement for constrained DNA ends preventing loop formation across long distances^63^. Importantly, because the 3’ junction is likely occupied by other proteins (e.g. the replisome during S phase), our data suggest that ssDNA looping by Dpb11 and RPA will still shorten the gap length between the 5’ junction and internal sites. Consequently, we suggest that Dpb11 bridges formed during checkpoint signaling may bring activators of Mec1 located at the gap junction and Mec1-Ddc2 at internal gap sites in closer proximity, allowing for colocalization and subsequent Mec1 activation. This may enable amplification of checkpoint signaling for longer gaps (**Figure** 5B-ii). Further studies are necessary to determine the exact physical interactions and protein composition that make up bridges formed in the presence of Dpb11 and RPA, both separately and together.

### Dpb11 directly positions Mec1 within the activation radius: recruiting Mec1-Ddc2 to ss-dsDNA junctions

Independent recruitment of Mec1-Ddc2 to RPA-ssDNA has been well established^6,8-11^ and reproduced in the current study. We present an additional pathway for Mec1-Ddc2 recruitment specifically to a gap junction, namely, via Dpb11. In the presence of excess Dpb11, Mec1-Ddc2 was significantly enriched at ss-dsDNA junctions compared to the theoretical model for non-specific RPA-ssDNA binding. Indeed, the distribution of Mec1-Ddc2 binding sites on RPA-coated gapped DNA in the presence of Dpb11 adopts the same pattern as Dpb11 on RPA-coated gapped DNA. This suggests that the interactions between Dpb11 and DNA junctions are driving the assembly of the Mec1-Ddc2/Dpb11 complex at a ss-dsDNA junction. We also report that colocalization events between Mec1-Ddc2 and Dpb11 at the junctions are predominantly the result of protein-protein interactions in solution or on DNA (initiated by Dpb11-junction binding). This is supported by previous studies demonstrating that interaction between Dpb11 and Mec1-Ddc2 are needed for a functional DNA damage checkpoint^14,26^. Although Dpb11 is evidently able to recruit Mec1-Ddc2 to either a 5’ or 3’ junction, we expect complete checkpoint activation (i.e. activation of both Mec1 and Rad53) to occur most frequently at a 5’ junction. We presume that, in a cellular context, replisome proteins, TLS polymerases, or other repair factors that occupy the 3’ junction may prevent Dpb11 recruitment to these sites, thereby increasing enrichment at the 5’ junction specifically. Therefore, targeting by Dpb11 directly positions the checkpoint kinase within the activation radius created by Dpb11 and/or 9-1-1 at the 5’ junction (**Figure** 5B-i).

In contrast to the current model for Dpb11 recruitment, our data indicate that Dpb11 can recognize and bind to gapped DNA in the absence of 9-1-1. While we propose that this facilitates Mec1 activation, it is important to distinguish between direct activation by the Dpb11 MAD versus indirect activation by recruitment, scaffolding, or other means. The current study does not rule out either possibility, and future work will be necessary to differentiate more explicitly. Important *in vivo* studies have demonstrated that, during S phase, robust activation of the replication stress checkpoint can be achieved without Ddc1^64,65^. We suggest that our observed recruitment of Mec1-Ddc2 to ssDNA gap junctions by Dpb11 may help to explain this 9-1-1-independent pathway. Because the Mec1/ATR activation pathway contains several redundancies, it is not entirely surprising that a Dpb11-only pathway may be able to compensate for a lack of 9-1-1.

Dpb11 is able to both bind and activate Mec1 in solution (without DNA or 9-1-1) in an ensemble biochemical assay^13^; the same is true for TopBP1 and ATR^66^. *In vivo*, however, checkpoint activation does not occur in the absence of DNA damage or replication stress. We speculate that even if the Mec1 kinase may be activated in solution, it still must be recruited to ssDNA in order to phosphorylate Rad53 (which interacts only weakly with Mec1 in solution^13^ but is sufficiently scaffolded by Dpb11/Rad9 interactions^31^) and trigger downstream steps (**Figure** 5B-iii). It is possible that stoichiometry, local concentrations, and changes in protein expression in response to genotoxic stress help to regulate these steps. Thus, we conclude that Dpb11 is able to directly recruit Mec1-Ddc2 to RPA-coated ssDNA gap junctions independently of 9-1-1 and propose a model in which this facilitates overall Mec1/checkpoint activation. It will also be necessary to incorporate 9-1-1 into future studies, as we and others have performed structural analysis on the checkpoint clamp and its loader^20,21,23^. Overall, our current data support a model that positions both 9-1-1 and Dpb11 at the 5’ junctions of RPA-coated gaps.

Taken together, our data support a model in which the combination of DNA gap recognition and scaffold building by Dpb11 facilitates Mec1 positioning within the activation radius. These findings significantly expand the role(s) of Dpb11 during the initial steps of the Mec1-mediated DNA damage checkpoint pathway.

## METHODS

### Protein expression and purification

*RPA*. Expression and purification of *Saccharomyces cerevisiae* (Sc) RPA was performed as described^67,72^.

#### Mec1 expression vector

The Sc Mec1 gene was synthesized by Biomatik. Mec1(a) contained a C-terminal PreScission protease site followed by a 3xFLAG tag. Mec1(b) contained a C-terminal S6 sequence^73^ followed by a 3xFLAG tag. Mec1(a) or Mec1(b) synthetic genes were inserted into pRS403 (His selection) and the sequences were confirmed by Sanger sequencing over the entire inserted region by GENEWIZ from Azenta Life Sciences. The Mec1(b) integration/expression plasmid has been deposited to Addgene (# 240737).

#### Ddc2 expression vector

The gene encoding Sc Ddc2 was obtained by PCR from genomic DNA isolated from Sc strain S288C. The PCR product was cloned into the pRS404 plasmid (Trp selection) and the sequence was confirmed by Sanger sequencing over the entire inserted region by GENEWIZ from Azenta Life Sciences. The Ddc2 integration/expression plasmid has been deposited to Addgene (# 240738).

#### Dpb11 expression vector

The Dpb11 gene (Sc strain S288C) was inserted into the pET-11a plasmid. The gene was then amplified using primers that incorporated a 3xFLAG tag at the C-terminus. The PCR product was cloned into the pRS405 plasmid (Leu selection) and the sequence was confirmed by Sanger sequencing by GENEWIZ from Azenta Life Sciences. The Dpb11 integration/expression plasmid has been deposited to Addgene (# 240739).

#### Mec1-Ddc2 expression and purification

The Mec1-pRS403 and Ddc2-pRS404 plasmids, encoding genes for over-expression under the GAL1/10 promotor, were integrated into the chromosome of yeast strain OY001 as described before for expression of other genes using the pRS402-pRS406 plasmid series^67^. Cells were initially grown under selection to make an overnight growth that was subsequently divided into 24 × 2L flasks (1L/flask) and grown to OD 600 at 30 °C before inducing with GAL as described^67^. After 6 h induction, cells were harvested and resuspended in a minimal volume of Buffer A (20 mM Tris-HCl pH 7.5, 10% glycerol, 1 mM EDTA, YPI) plus 1 M NaCl. The yeast protease inhibitor cocktail (YPI) was used at a 1:1,000 dilution. Cells were then dripped into liquid nitrogen, producing beads of frozen cells that were stored at -80 °C. Frozen cells were then lysed simultaneously using two freezer mills (SPEX SamplePrep) with the following grinding protocol: 15 cycles (2 min grinding followed by 5 min chill in liquid nitrogen) at rate 9. The final ground material was resuspended in 30 ml Buffer A + 1 M NaCl. The mixture was slowly agitated until fully unfrozen at 4 °C. Cell debris was removed in a 1 h spin at 12,500 rpm at 4 °C in a Sorvall Lynn 6000 centrifuge and F14-6-250 rotor.

For Mec1(a)-Ddc2, the 100 mL supernatant was collected and 1 mL FLAG-Ab beads were added, followed by gentle agitation for 1 h at 4 °C. The beads were pelleted using a Sorvall BP8 centrifuge rotor at 1,800 rpm for 9 min at 4 °C. The pelleted beads were washed three times at 4 °C. For each wash, the pelleted beads were resuspended with 100 ml Buffer A + 1 M NaCl + 1 mM ATP + 4 mM MgCl_2_, gently agitated for 15 min, and collected by centrifugation. The pelleted beads were then resuspended in a total of 10 ml Buffer A + 1 M NaCl, then packed into a C-column (GE Healthcare) that was pre-equilibrated with Buffer ET (20 mM Tris-HCl pH 7.5, 10% glycerol, 1 mM EDTA, YPI, 300 mM NaCl). The column was washed using Buffer ET and then eluted in Buffer ET + 0.2 mg/ml FLAG peptide using a 0.6 mL/min flow rate. Fractions were analyzed for Mec1-Ddc2 by SDS-PAGE, and peak fractions were aliquoted, snap frozen in liquid nitrogen, and stored at -80 °C (yield 0.3 mg). Protein concentration was estimated using Bradford Protein Stain (Bio-Rad Labs) using BSA as a standard. Mec1(b)-Ddc2 was prepared exactly as above, except that Buffer ET was replaced with Buffer EH (50 mM HEPES pH 7.5, 10% glycerol, 0.01% NP40, YPI, 300 mM KCl).

#### Dpb11 expression and purification

The pRS405 plasmid encoding Dpb11 under control of the GAL1/10 promotor was integrated into yeast strain OY001 and cells were grown and prepared as described above. Briefly, 24 L yeast cells were grown then “drip frozen” in liquid nitrogen. Cells were lysed using two simultaneous freezer mills (SPEX SamplePrep) using the same protocol as above. The final ground material was resuspended in 80 ml Buffer A + 800 mM NaCl and thawed by gentle stirring 4 °C. Cell debris was removed by centrifugation for 1 h at 12,500 rpm at 4 °C in a Sorvall Lynn 6000 centrifuge and F14-6-250 rotor. Then, 1 mL of FLAG-Ab beads were added to the supernatant, followed by gentle agitation for 1 h at 4 °C. The beads were washed 4 times with Buffer A + 800 mM NaCl. The final washed beads were packed into a C-column and the column was eluted with 0.2 mg/ml 3xFLAG peptide and fractions were analyzed by SDS-PAGE. Peak fractions were pooled and then concentrated using a 12 mL Amicon Ultra 30K centrifugal filter (yield 1.34 mg). Fractions were aliquoted, snap frozen in liquid nitrogen, and stored at -80 °C.

### N-terminal amine labeling with NHS-ester-functionalized fluorophores

#### Fluorescent Dpb11

Purified Dpb11 was first dialyzed into Buffer B (50 mM HEPES-KOH pH 7, 500 mM NaCl, 1 mM DTT, 1 mM EDTA, 10% glycerol). Dpb11^AF555^ was then prepared by incubating Dpb11 with a 5-fold molar excess of Alexa Fluor 555 (AF555) NHS (Life Technologies) for 1.5 hours at 4 °C. The reaction was stopped by adding Tris-Acetate pH 7.5 to a final concentration of 25 mM.

Dpb11^LD655^ was prepared by incubating dialyzed Dpb11 with a 5-fold molar excess of LD655 NHS (Lumidyne Technologies) for 1.5 hours at 4 °C. The reaction was stopped by adding Tris-Acetate pH 7.5 to a final concentration of 25 mM. Excess free dye was then removed via Zeba Dye and Biotin Removal Spin Columns pre-equilibrated with Buffer C (20 mM Tris-HCl pH 7.5, 500 mM NaCl, 12% glycerol) according to the manufacturer protocol. 1 mM TCEP and 0.25 mM EDTA were added to the washed, labeled protein.

Fluorescently labeled proteins were validated by SDS-PAGE (**Figure** S1A). Fluorescent protein bands were visualized using a laser scanner for fluorescence with appropriate excitation and emission filters (Typhoon, Amersham) and total protein content was stained with Quick Blue protein stain (IBI Scientific). Proteins were then aliquoted, flash frozen in liquid nitrogen, and stored in the dark at -80 °C.

#### Fluorescent Mec1-Ddc2

Mec1-Ddc2 was dialyzed into Buffer D (50 mM HEPES-KOH pH 7, 300 mM NaCl, 1 mM DTT, 0.5 mM EDTA, 0.4 mM PMSF, 10% glycerol). Mec1-Ddc2^AF647^ was then prepared by incubating dialyzed Mec1(a)-Ddc2 with a 5-fold molar excess of Alexa Fluor 647 NHS (Life Technologies) for 1 hour at 4 °C. The reaction was stopped by adding Tris-Acetate pH 7.5 to a final concentration of 25 mM. Excess free dye was removed by three consecutive wash steps over an Amicon Ultra 50K centrifugal filter in Buffer E (20 mM Tris-Acetate pH 7.5, 300 mM NaCl, 1 mM DTT, 0.5 mM EDTA, 15% glycerol, 0.04 mg/ml BSA).

Mec1-Ddc2^LD555^ and Mec1-Ddc2^LD655^ were prepared by incubating Mec1(b)-Ddc2 with a 5-fold molar excess of LD555 NHS or LD655 NHS (Lumidyne Technologies), respectively, for 10 minutes at 4 °C. The reaction was stopped by adding Tris-Acetate pH 7.5 to a final concentration of 25 mM. Excess free dye was then removed via Zeba Dye and Biotin Removal Spin Columns pre-equilibrated with Buffer F (20 mM Tris-HCl pH 7.5, 300 mM NaCl, 15% glycerol) according to the manufacturer protocol. 1 mM TCEP and 0.25 mM EDTA were added to the washed, labeled protein. Preparations of Mec1-Ddc2^AF647^, Mec1-Ddc2^LD555^, and Mec1-Ddc2^LD655^ were validated and quantified as above (**Figure** S1B), then aliquoted, flash frozen, and stored at -80 °C.

### Ensemble phosphorylation assay

Phosphorylation reactions (20 μl) were performed in SM Buffer (25 mM Tris-HCl pH 7.5, 100 mM NaCl, 10 mM Mg(CH_3_COO)_2_, 2 mM TCEP, 0.1 mM EDTA, 0.04 mg/ml BSA, 1% glycerol) with 200 nM RPA and 1 nM 18-primed ΦX174 ssDNA. RPA/DNA mixes were pre-incubated at 30 °C for 1 minute before adding 0 or 10 nM Dpb11 and incubating for another 5 minutes at 30 °C. Reactions were initiated with 10 nM Mec1-Ddc2, 100 μM cold ATP, and 75 nM (4.5 μCi) γ-^32^P ATP. After the indicated times at 30 °C, reactions were stopped with 5 μl of 5X SDS gel loading buffer. Reaction products were separated by SDS-PAGE on a 4-20% Tris-Glycine gel and stained with Quick Blue (IBI Scientific). The gel was then exposed to a phosphor imaging screen and scanned on an Amersham Typhoon. This assay was repeated with all three fluorescently labeled preps of Mec1-Ddc2 used in this study (Mec1-Ddc2^AF647^, Mec1-Ddc2^LD555^, and Mec1-Ddc2^LD655^).

### Preparation of DNA substrates with site-specific nicks for single-molecule experiments

The DNA substrate was prepared as previously described^74^. A 3 kb AT-rich fragment was PCR-amplified from plasmid pLW58 (based on pAS1^68^) using primers PriF and PriR (**Key Resources Table**). Each primer introduced an Nb.BbvCI nicking site as well as an NheI (PriF) or BsrGI (PriR) cloning restriction site. The 3 kb PCR product was cloned into the 15.9 kb plasmid pRGEB32^69^ via NheI/BsrGI (PCR product) and NheI/BsiWI (pRGEB32) to generate pJF3KB1.

Purified pJF3KB1 was linearized by restriction digest with BsaI-HF v2 (all restriction enzymes from New England Biolabs) and the resulting 20 bp fragment was removed by PEG precipitation. Biotinylated oligo pairs (Oli1+Oli2, Oli3+Oli4) were annealed and ligated (T4 DNA ligase, New England Biolabs) to the two BsaI overhangs of the linearized plasmid. Ligase was then heat inactivated. The DNA was nicked on the same strand at the two Nb.BbvCI nicking sites at either end of the 3 kb AT-rich region. The endonuclease was then heat inactivated and the DNA was purified away from excess oligos by PEG precipitation. Aliquots of the intermediate linearized pJF3KB1 and the final biotinylated/nicked substrate were digested with SphI and SpeI to check for successful BsaI restriction, PEG purification, and introduction of the oligo pairs at both ends of the DNA. The substrate was stored in TE buffer at 4 °C. All synthetic oligonucleotides were ordered from Integrated DNA Technologies (**Key Resources Table**).

### Generation of 3 kb ssDNA gaps for single-molecule experiments

Gapped DNA was generated in situ on a C-Trap instrument^42^ (Lumicks) at room temperature. All buffers were filtered through 0.22 μm PVDF Durapore membranes. Under constant flow, single 3.13 or 4.38 μm streptavidin-coated polystyrene microspheres (Spherotech) diluted in PBS were caught in each of two optical traps in the first channel of a laminar flow cell (Lumicks). Trapped beads were then moved into a second channel containing 19 kb end-biotinylated nicked DNA (prepared as above) in PBS under flow. DNA ends were caught by beads via biotin/streptavidin interaction. Force-extension curves of each tethered DNA were collected and compared to the expected worm-like chain (WLC) model (below) to confirm that only one DNA molecule of the expected length was attached between the two beads. Finally, trapped beads (with tethered DNA) were moved into a third channel containing no-salt buffer (20 mM Tris pH 7.4). Under flow, force-induced DNA melting was achieved by moving beads apart, causing complete melting and removal of the AT-rich fragment between the two nicks. Flow was turned off and force-extension curves of the now gapped DNA were collected and compared to the expected worm-like chain plus freely jointed chain (WLC-FJC) hybrid model (below) to confirm the desired DNA structure (**Figure** S2A-C).

### Fitting of DNA force-extension curves

All experimental DNA force-extension data were analyzed according to appropriate polymer chain models^75,76^. The extension distance (*D*, nm) of duplex DNA can be modeled as a function of force (*F*, pN) with a worm-like chain (WLC) model^77-79^, approximated as:

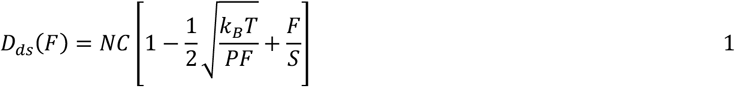

where *N* is the DNA length (# residues), *C* is the contour length of one DNA residue (nm/residue), *k*_*B*_*T* is the Boltzmann constant (4.114 pN × nm), *P* is the persistence length (nm), and *S* is the stretch modulus (pN). Single-stranded DNA is modeled using a freely jointed chain (FJC) model^80^:

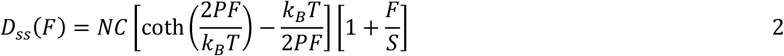

Combining Equations 1 and 2 allows us to consider DNA which contains both double-stranded and single-stranded regions, such as our gapped DNA substrate:

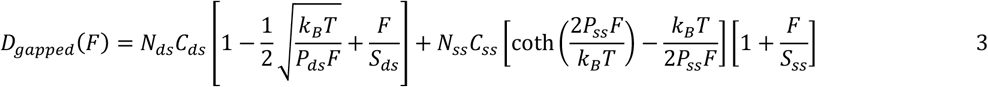

Because polymer behavior of DNA is significantly affected by experimental conditions such as ionic strength and pH, relevant parameters (contour length, persistence length, and stretch modulus) for dsDNA, ssDNA, and RPA-ssDNA were determined empirically in SM Buffer (25 mM Tris-HCl pH 7.5, 100 mM NaCl, 10 mM Mg(CH_3_COO)_2_, 2 mM TCEP, 0.1 mM EDTA, 0.04 mg/ml BSA, 1% glycerol) at room temperature under no flow. Using Bluelake software on a C-Trap instrument, force-extension curves were first obtained for the nicked DNA substrate (**Figure** S7A) to model dsDNA. DNA was stretched from 2.5 μm to 6.5 μm by slowly moving one trap/bead. Force was measured on the stationary trap and downsampled to 50 Hz. All data collected between ∼1 pN and 30 pN were plotted in MATLAB and globally fit to Equation 1 (*N* = 18.851 kb) using least squares nonlinear regression. Force-extension data were then collected as above for gapped DNA (**Figure** S7B). These data were fit to Equation 3 using parameters we obtained above for dsDNA (*C*_*ds*_ = 0.326 nm/bp, *P*_*ds*_ = 36.578 nm, *S*_*ds*_ *=* 1132.1 pN) to determine parameter values pertaining to the single-stranded region. Finally, force-extension data were collected in the presence of 100 nM RPA (**Figure** S7C) and fit to Equation 3 to determine parameter values pertaining to the single-stranded region bound by RPA.

### Single-molecule imaging

Using the U-Flux microfluidics system (Lumicks), the laminar flow cell was passivated by flowing in 1 mg/ml BSA (New England Biolabs) in PBS for 15 minutes followed by 0.5% w/v Pluronic F127 (Sigma-Aldrich) in PBS for 15 minutes. The flow cell was then equilibrated with appropriate buffers for each channel.

All kymographs and protein-DNA binding position were collected on a C-Trap with Bluelake software (Lumicks) under no flow at room temperature. Beads were held at a constant distance and all data was collected under low force (<20 pN) on DNA. For each kymograph, gapped DNA (prepared as above) was first brought into a channel of the laminar flow cell containing the dsDNA stain YO-PRO-1 (ThermoFisher Scientific) in PBS to confirm location of gap junctions. The DNA was then brought into a channel containing proteins as specified for each experiment in SM Buffer with oxygen scavenging reagents (2.5 mM protocatechuic acid, 25 nM protocatechuate-3,4-dioxygenase, 1 mM cyclooctatetraene, 1 mM nitrobenzyl alcohol, 1 mM Trolox). YO-PRO-1 readily dissociates from DNA and was not present during protein binding. Dpb11 reactions included 10-50 nM Dpb11^AF555^ or Dpb11^LD655^ supplemented with 150 nM unlabeled Dpb11. Dpb11 + RPA reactions included 5-30 nM Dpb11^AF555^ and 100 nM unlabeled RPA. Mec1-Ddc2 reactions included 30 nM Mec1-Ddc2^AF647^, Mec1-Ddc2^LD655^ or Mec1-Ddc2^LD555^, and 100 nM unlabeled RPA. Mec1-Ddc2 + Dpb11 reactions included 30 nM Mec1-Ddc2^AF647^ or Mec1-Ddc2^LD555^, 200 nM unlabeled Dpb11, and 100 nM unlabeled RPA. Mec1-Ddc2 + Dpb11 dual-color experiments were performed with 10 nM Mec1-Ddc2^LD555^, 20 nM Dpb11^LD655^, 180 nM unlabeled Dpb11, and 100 nM unlabeled RPA. Kymographs were collected at a frame rate of 0.02 – 0.1 seconds/line and a pixel size of 50 or 100 nm.

### Kymograph processing and analysis

The photon count file of each kymograph was obtained using the lumicks.pylake Python library and a custom GUI Python script titled “C-Trap.h5 File Visualization GUI” (retrieved from https://harbor.lumicks.com/)^70^. Image files were processed using Fiji (NIH)^71^. Initial binding positions of fluorescent proteins on DNA were determined using a one-dimensional Gaussian fit to the fluorescence intensity of the kymograph. Position data were converted from pixels to nm, then converted from nm to nt based on the appropriate WLC/FJC model (above), using the known DNA extension/force and YO-PRO-1 staining as a marker for duplex regions. Fluorescent proteins on DNA were identified in kymographs and categorized as stationary or mobile (if position changed more than 150 nm during its duration on DNA).

Histograms of protein binding positions on 3 kb-gapped DNA were generated using bins (width 506 nt) centered at the 5’ and 3’ gap junctions and distributed evenly throughout the 3,036 nt gap. A single histogram bin was used for dsDNA on either side of the gap, normalized to the appropriate length, because only a single binding event was observed on dsDNA for all experiments/conditions. Protein binding distributions were compared against the theoretical distribution for non-specific ssDNA-binding by χ^2^ test for goodness of fit. The model for non-specific ssDNA-binding was generated based on two assumptions: (1) no sites on dsDNA are occupied, and (2) every site on ssDNA is equally likely to be occupied. At ss-dsDNA junctions, histogram bins include one junction site plus equal amounts of both dsDNA and ssDNA. Thus, the model predicts half of the binding frequency for a junction bin (i.e. 253 nt ssDNA) compared to a bin with only ssDNA (i.e. 506 nt ssDNA).

### Single-molecule DNA pulling experiments

All force-extension data for DNA looping and bridging experiments were collected on a C-Trap with Bluelake software under no flow at room temperature. The flow cell was prepared and passivated as described above. 18.851 kb DNA (nicked or gapped), tethered between two optically trapped beads, was brought into a channel containing proteins as specified for each experiment in SM Buffer with oxygen scavenging reagents. Dpb11 reactions contained 200 nM unlabeled Dpb11. RPA reactions contained 100 nM unlabeled RPA. Dpb11+RPA reactions contained 200 nM Dpb11 and 100 nM RPA. Beads were brought into close proximity (1.5 – 2.5 μm apart) and held for 10 seconds to allow for tethered DNA to encounter itself according to the polymer chain in three dimensions under no tension, forming loops. DNA was extended by moving one optically trapped bead 0.8 μm/sec until the force reached 20-30 pN and was held at this extended position for 10 seconds. The DNA was then relaxed by moving the optically trapped bead back in the opposite direction to confirm that all bridges were disrupted and the DNA behaved as the unmodified, full-length conformation. To prevent scoring repeated data on the same protein bridges, pulling tests (i.e. full cycle of extension and relaxation) were repeated on the same DNA molecule no more than 5 times, and were only counted if all bridges were confirmed to be lost (by a negative pulling test) between pulls.

The corresponding force, extension, and time data for each DNA was obtained using the lumicks.pylake Python library and the custom GUI Python script titled “C-Trap.h5 File Visualization GUI”. Each individual pull (i.e. a single passage from completely relaxed to maximum extension) was processed in Microsoft Excel. Protein bridge breaks were identified manually by looking for drops in force with increasing DNA extension and force-extension curves were separated based on these breaks. Data were then fit to the WLC-FJC model for gapped DNA (Equation 3) to solve for the length of the ssDNA or RPA-ssDNA region (*N*_*ss*_) using the empirical parameters determined above. *N*_*SS*_ at maximum DNA extension was checked against the full-length model and assumed to be 3.036 kb. The break force is defined as the maximum force measured before breaking. The loop length is defined as the difference in DNA length before and after a break (*N*_*ss*_^*post-break*^ – *N*_*ss*_^*pre-break*^).

### Mass photometry

Stoichiometry data were collected with a OneMP mass photometer using AcquireMP software v2.4.0 (Refeyn). Protein mixes were prepared as specified for each experiment in MP Buffer (25 mM Tris-HCl pH 7.5, 250 mM NaCl, 10 mM Mg(CH_3_COO)_2_, 2 mM TCEP, 0.1 mM EDTA, 4% glycerol) and applied to a prepared glass coverslip. For each sample, light scattering signal was obtained from a 60 second movie (100 frames/second) and particle masses were calculated using a calibration curve of known protein standards (bovine serum albumin, 66 kDa; beta amylase, 224 kDa; thyroglobulin, 670 kDa). Histograms of particle masses were plotted and Gaussians were fit to the data by nonlinear regression in MATLAB (MathWorks).

## ACKNOWLEDGEMENTS

The authors thank Jeff Finkelstein for help with cloning, and Olga Yurieva and Dan Zhang for protein expression and purification. We thank Noa Dahan for help with mass photometry. Plasmid pLW58 was a gift from Ling Wang. pRGEB32 was a gift from Yinong Yang (Addgene plasmid # 63142). John Watters provided custom python scripts. Nina Yao provided 18-primed Φx174 DNA. This work was supported by the National Institute of Health (R01GM149862 to SL and R35GM148159 to MEO), the Howard Hughes Medical Institute (to MEO), the Breast Cancer Research Foundation (BCRF-24-068 to MEO), the Marlene Hess Center for Research on Women’s Health and Biomedicine at Rockefeller University (to SL), and an Anderson Center for Cancer Research postdoctoral fellowship (to ECB).

## AUTHOR CONTRIBUTIONS

ECB conceived the research under the guidance of MEO and SL. ECB performed and analyzed all experiments. ECB wrote the manuscript, which was reviewed, discussed, and edited by MEO, SL, and GC.

