## Supplemental figures for "Dpb11 facilitates the colocalization of Mec1-Ddc2 with its activators on gapped DNA"

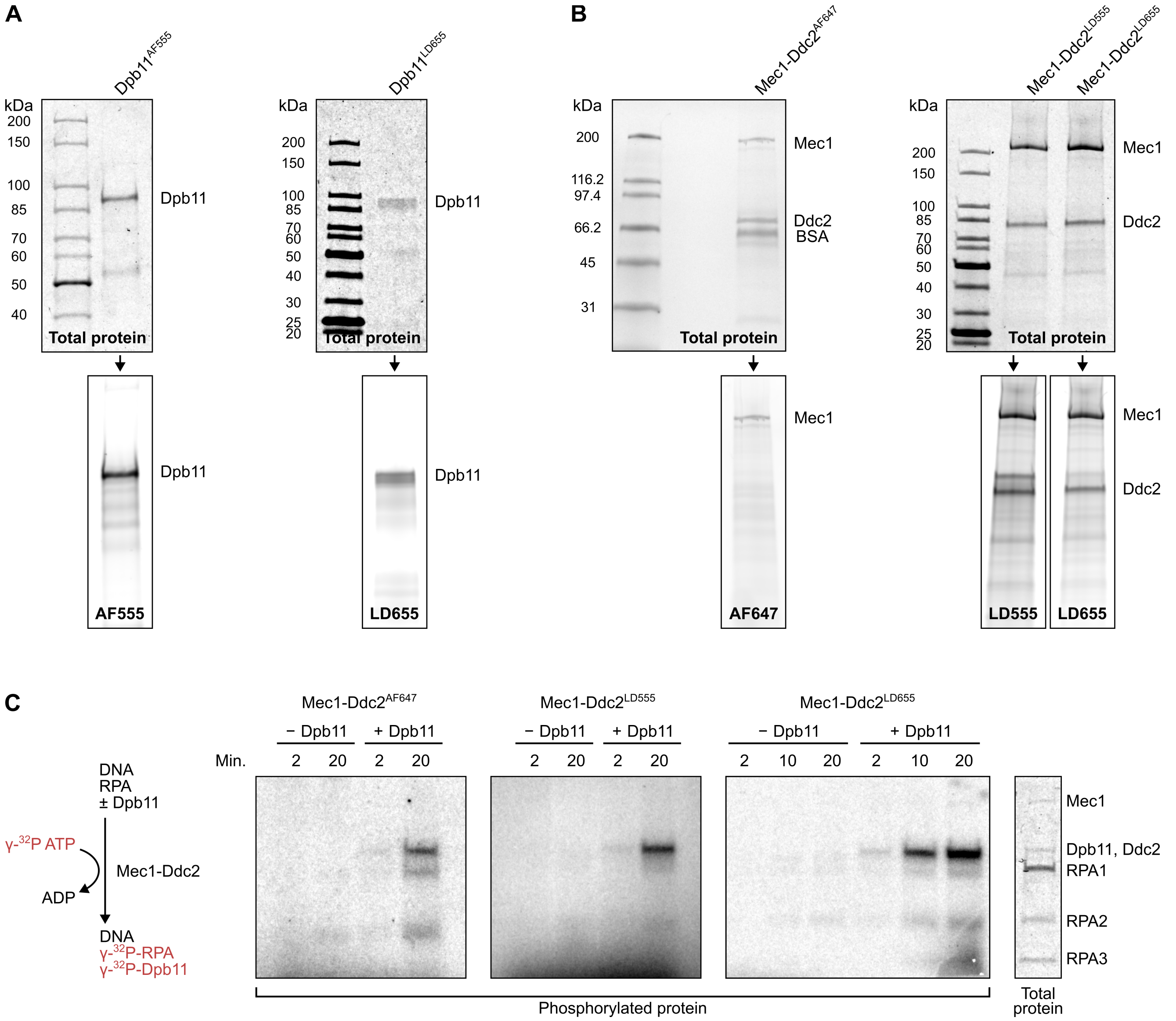


**Figure S1. Proteins used in this study.**

(A) SDS-PAGE of purified Dpb11$^{AF555}$ and Dpb11$^{LD655}$, stained for total protein (top) and scanned for fluorescence (bottom). (B) SDS-PAGE of purified Mec1-Ddc2$^{AF647}$, Mec1-Ddc2$^{LD555}$, and Mec1-Ddc2$^{LD655}$, stained for total protein (top) and scanned for fluorescence (bottom). (C) Phosphorylation assay to test Dpb11 activation of Mec1 kinase. Purified and fluorescently labeled Mec1-Ddc2 was incubated with ATP, 18-primed ΦX174 ssDNA, and RPA, with or without Dpb11 for the indicated times. Phosphorylation reactions were separated by SDS-PAGE, stained for total protein (right), and phosphorimaged to detect incorporation of γ-$^{32}$P ATP.

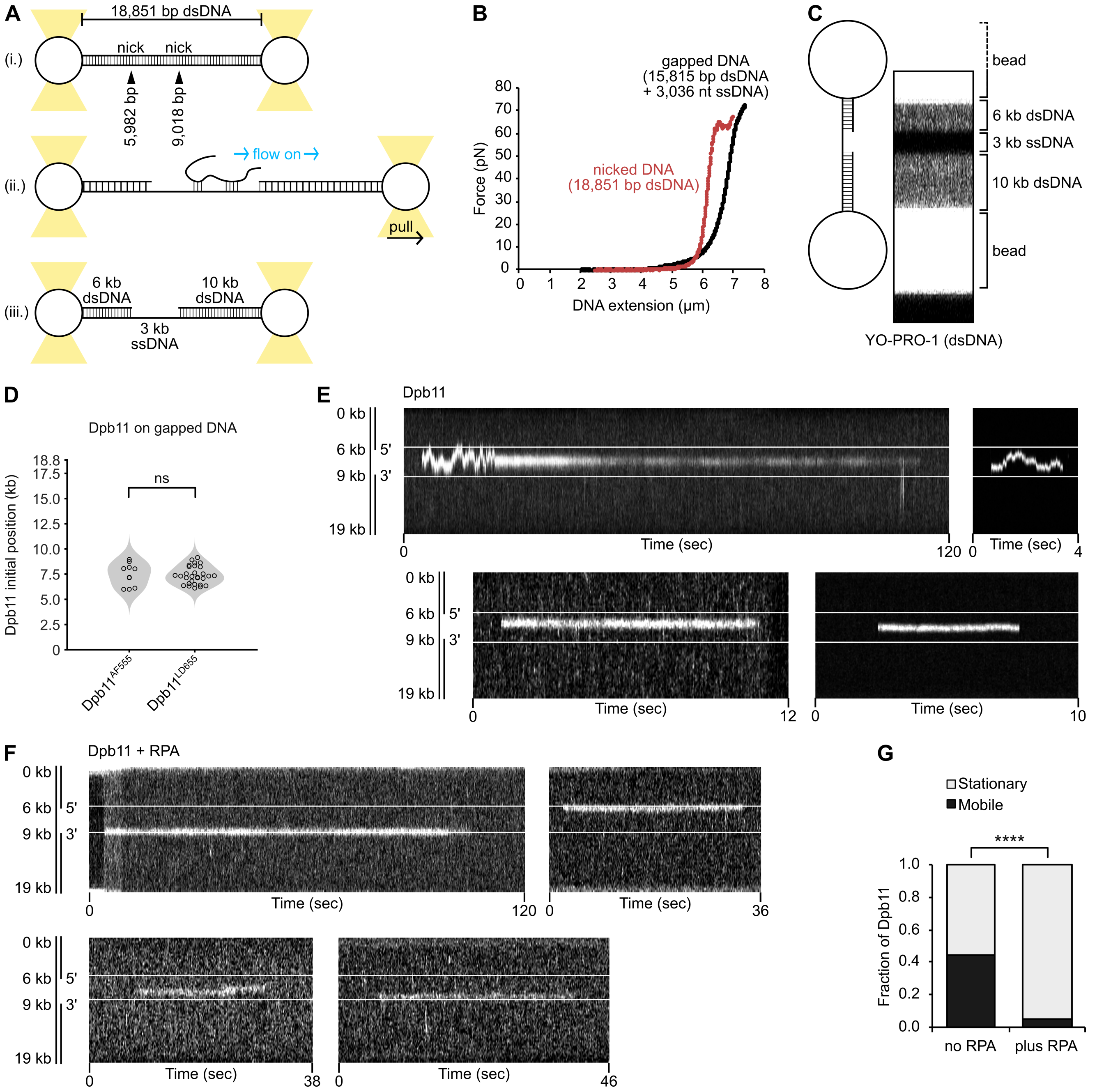


**Figure S2. Single-molecule imaging of Dpb11 on gapped DNA, related to Figure 1.**

(A) Generation of 3 kb-gapped DNA for single-molecule experiments. (B) Sample experimental data confirming the change in force-extension curves before (red) and after (black) generation of the ssDNA gap. (C) Kymograph (15 sec) showing YO-PRO-1 staining of dsDNA on 3 kb-gapped DNA substrate. (D) Comparison of DNA-binding behavior of Dpb11 labeled with AF555 and LD655. Samples were compared by Mann Whitney U test. ns, not significant. (E and F) Additional kymographs of $Dpb11^{AF555}$ and $Dpb11^{LD655}$ on 19 kb DNA with 3 kb single-stranded gap (6–9 kb) in the absence (E) or presence (F) of RPA. See **Figures** 1B and 1D, respectively. (G) Fraction of stationary and mobile Dpb11 particles on 3 kb-gapped DNA in the absence or presence of RPA. **** $p = 6.8\times10^{-5}$ by $\chi^2$ test for independence.

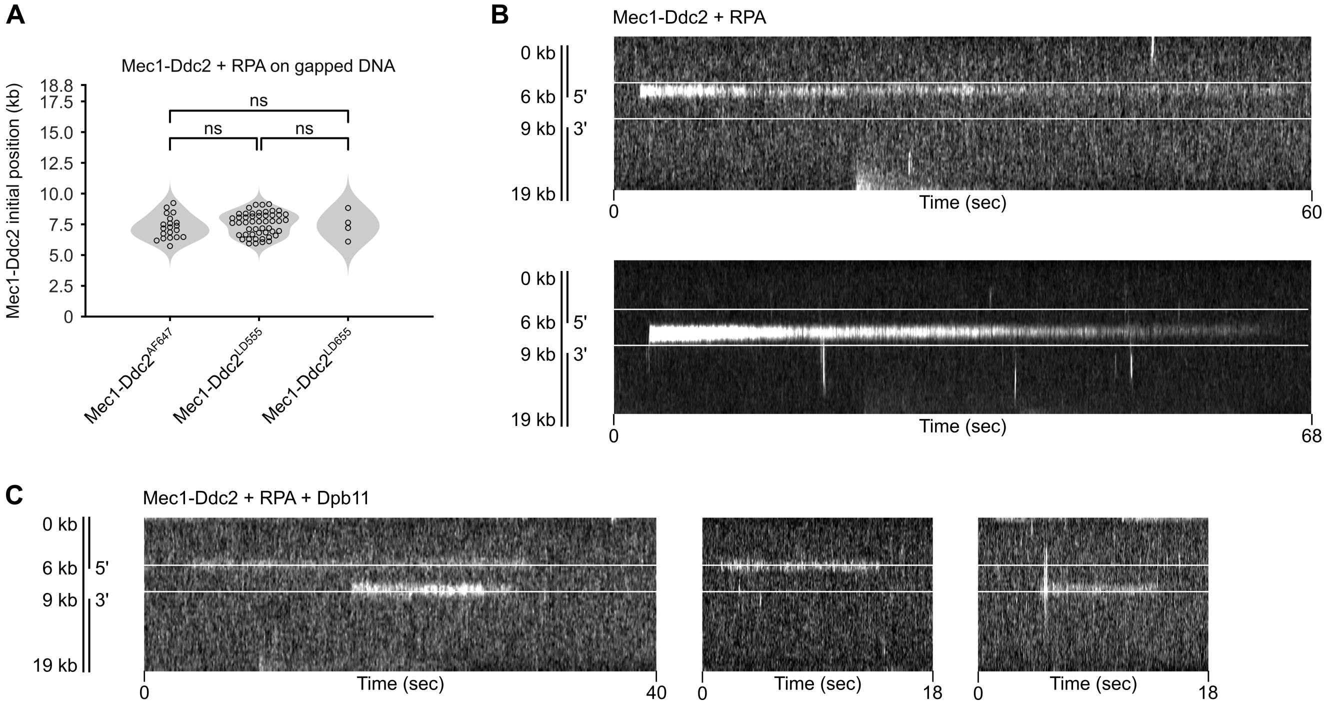


**Figure S3. Single-molecule imaging of Mec1-Ddc2 on gapped DNA, related to Figure 2.**

(A) Comparison of DNA-binding behavior of Mec1-Ddc2 labeled by AF647, LD555, and LD655. Samples were compared by Mann Whitney U test. ns, not significant. (B and C) Additional kymographs of Mec1-Ddc2$^{AF647}$, Mec1-Ddc2$^{LD655}$, and Mec1-Ddc2$^{LD555}$ with unlabeled RPA on 19 kb DNA with 3 kb single-stranded gap (6–9 kb) in the absence (B) or presence (C) of Dpb11. See **Figures** 2A and 2C, respectively.

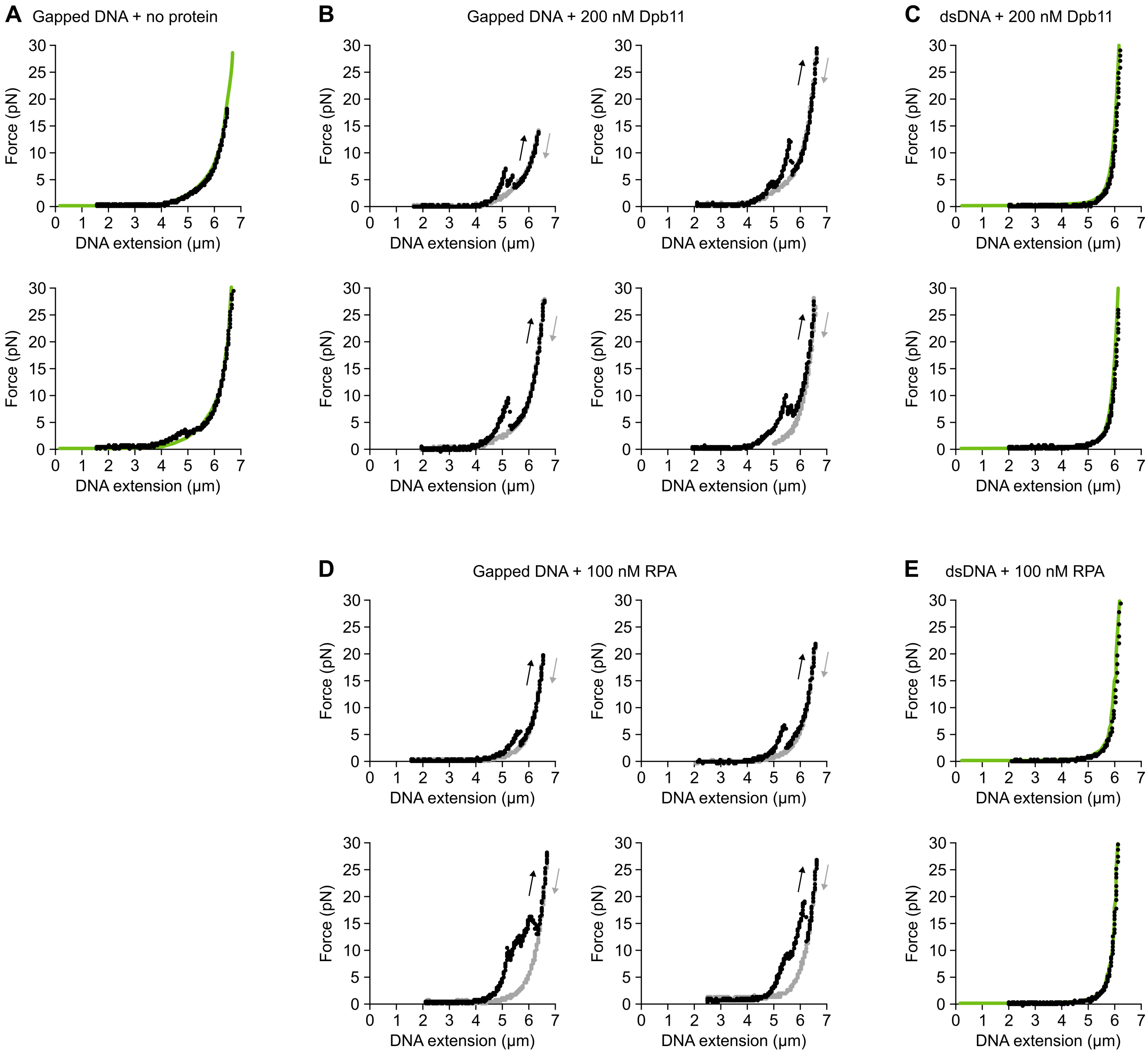


**Figure S4. DNA pulling tests, related to Figure 3.**

(A) Example DNA pulling tests of gapped DNA (15.815 kb dsDNA + 3.036 kb ssDNA) in the absence of proteins. Black circles are experimental data. Green line is the predicted hybrid worm-like chain (WLC) + freely-jointed chain (FJC) model (**Figure** S7B). (B) Additional example DNA pulling tests of gapped DNA in the presence of Dpb11. Black circles are stretching data and gray circles are relaxation data. See **Figure** 3B. (C) Example DNA pull tests of nicked DNA (18.851 kb dsDNA) in the presence of Dpb11. Black circles are experimental data. Green line is the predicted WLC model (**Figure** S7A). (D) Example DNA pull tests of gapped DNA in the presence of RPA. See **Figure** 3E. (E) Example DNA pull tests of nicked DNA in the presence of RPA. Black circles are experimental data. Green line is the predicted WLC model (**Figure** S7A).

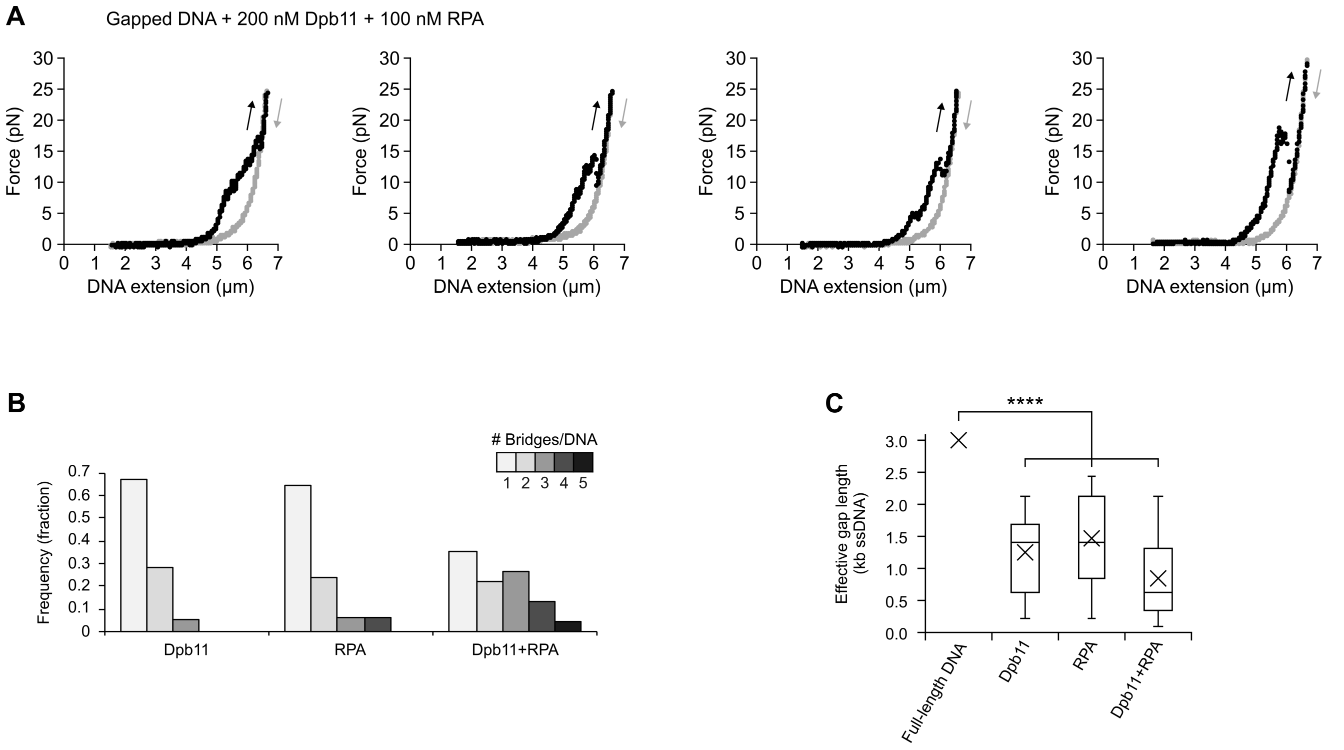


**Figure S5. Dpb11 bridges gapped DNA in the presence of RPA, related to Figure 3.**

(A) Additional example DNA pulling tests of 3 kb-gapped DNA in the presence of Dpb11 + RPA. Black circles are stretching data and gray circles are relaxation data. See **Figure** 3H. (B) Histograms showing the number of bridges per DNA molecule in the presence of Dpb11 only, RPA only, or both Dpb11+RPA. (C) The effective gap length (3.036 kb – sum of all loop lengths) in the presence of Dpb11 only, RPA only, or both Dpb11+RPA. **** $p < 1.0\times10^{-7}$ by one sample t-tests for all three conditions compared to full-length gapped DNA (3.036 kb ssDNA).

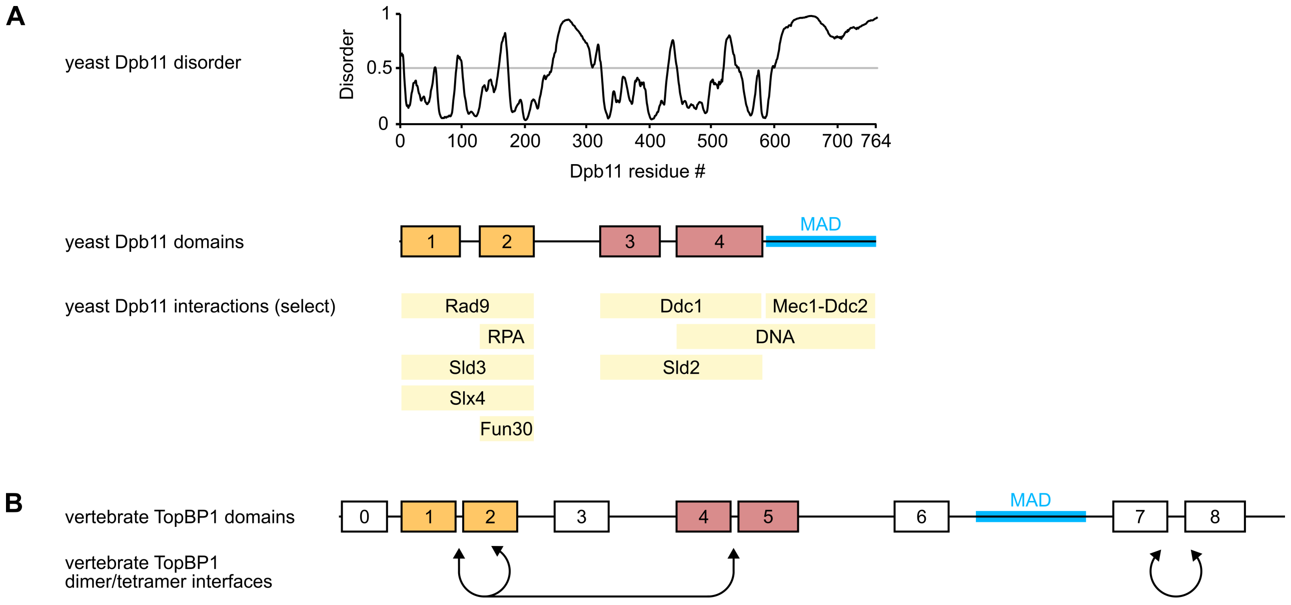


**Figure S6. Dpb11 domains, interaction partners, and homology, related to Figure 4.**

(A) Disorder predictions for yeast Dpb11 were obtained via the PONDR VSL2 algorithm[1]. BRCA1 C-terminal (BRCT) domains (numbered rectangles) and the Mec1-activating domain (MAD, blue) are shown below the disorder predictions. Select Dpb11 binding partners are displayed according to their respective interaction sites on Dpb11. Interactions involving DNA damage checkpoint activation: Rad9[2], Ddc1[3,4], Mec1-Ddc2[2,5,6]. Interactions with DNA and RPA, reported previously[7,8] and further investigated in this study. Interactions involved in other pathways: Sld2 and Sld3[9-12], Slx4[13,14], Fun30[15]. (B) Vertebrate TopBP1 BRCT domains (numbered rectangles) and Mec1/ATR activating domain (MAD, blue) are shown. Domains are colored according to their homologous counterparts in yeast. Self-oligomerization interfaces are indicated below domains[16].

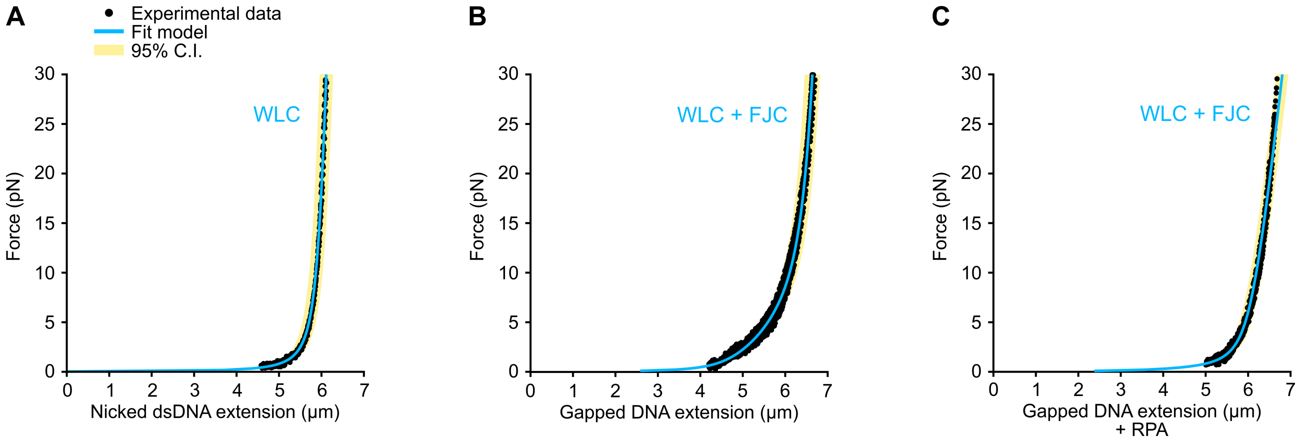


**Figure S7. DNA polymer modeling.**

Force-extension data for 18.851 kb DNA substrates in the absence of proteins were globally fit by least squares nonlinear regression to the worm-like chain model (WLC) or hybrid worm-like chain and freely-jointed chain model (WLC + FJC). Black circles: experimental data. Blue line: fit model. Yellow shading: 95% confidence interval of fit. (A) Nicked dsDNA (18,851 bp dsDNA) fit to WLC model (n = 4 DNA molecules). (B) Gapped DNA (15,815 bp dsDNA + 3,036 nt ssDNA) fit to WLC+FJC model (n = 6 DNA molecules). (C) Gapped DNA (15,815 bp dsDNA + 3,036 nt ssDNA) in the presence of RPA fit to WLC+FJC model (n = 6 DNA molecules).
